# Trade-offs in provisioning and stability of multiple ecosystem services in agroecosystems

**DOI:** 10.1101/350967

**Authors:** Daniel Montoya, Bart Haegeman, Sabrina Gaba, Claire de Mazancourt, Vincent Bretagnolle, Michel Loreau

## Abstract

Changes in land use generate trade-offs in the delivery of ecosystem services in agricultural landscapes. However, we know little about how the stability of ecosystem services responds to landscape composition, and what ecological mechanisms underlie these trade-offs. Here, we develop a model to investigate the dynamics of three ecosystem services in intensively-managed agroecosystems, i.e. pollination-independent crop yield, crop pollination, and biodiversity. Our model reveals trade-offs and synergies imposed by landscape composition that affect not only the magnitude but also the stability of ecosystem service delivery. Trade-offs involving crop pollination are strongly affected by the degree to which crops depend on pollination and by their relative requirement for pollinator densities. We show conditions for crop production to increase with biodiversity and decreasing crop area, reconciling farmers’ profitability and biodiversity conservation. Our results further suggest that, for pollination-dependent crops, management strategies that focus on maximising yield will often overlook its stability. Given that agriculture has become more pollination-dependent over time, it is essential to understand the mechanisms driving these trade-offs to ensure food security.

## Introduction

Human population growth and changes in diet preferences worldwide are generating a huge demand for food (Godfray et al 2010). To fulfil this increasing demand, agricultural intensification targets high crop yields. The merits of this approach are clear: the world annual production of cereals, grains, roots, tubers, pulses and oil crops has more than doubled, and the proportion of undernourished people in the world has decreased from 26% to 14% over the past 50 years (FAO 2009, 2011). But yields are no longer increasing in many major crops (Ray et al 2012) and show saturating responses to pesticide levels (Gaba et al 2016, Lechenet et al 2017), which suggests that the benefits of agricultural intensification have plateaued. Furthermore, these benefits have come at a considerable cost to biodiversity. This is particularly worrying for crops whose yield depends on ecosystem functions and services, such as pollination, whose provision has not traditionally been part of management policies (Pywell et al 2015, Tamburini et al 2016).

Global agriculture largely depends on animal pollination. It is estimated that 70% of 1,330 tropical crops (Roubik 2015) and 85% of crops in Europe (Williams 1994) benefit from animal pollination, and that pollinators can increase the production of 75% of the 115 most important crops worldwide (Aizen et al 2009). Although the three major crops in terms of biomass are independent of animal pollination (wheat, rice, corn), the cultivated area of pollination-dependent crops is expanding faster than the area of pollinator-independent crops (Breeze et al 2014, Aizen and Harder 2009). In contrast to the global increase in pollination-dependent agriculture, abundance and diversity of wild pollinators are declining worldwide (Goulson et al 2015). Honeybee (and sometimes bumblebee) colonies are used to substitute wild pollinator communities, yet the pollination services of wild pollinators cannot be compensated by managed bees because (i) pollinator-dependent crop land grows more rapidly than the stock of, e.g., honeybee colonies (Aizen et al 2009), (ii) wild insects usually pollinate crops more efficiently than honeybees (Garibaldi et al 2013), and (iii) honeybees may depress wild pollinator densities (Lindström et al 2016). Wild pollinators thus remain fundamental for agricultural pollination. In agricultural landscapes, the loss of semi-natural habitat is considered to be the first cause of wild pollinator declines (Kennedy et al 2013, Bretagnolle & Gaba 2015), as semi-natural elements (e.g. hedgerows, low-managed grasslands, forest patches) provide foraging, nesting and refuge habitats for pollinator communities (Kremen et al 2004). This land use change therefore leads to a continuous decrease of wild pollinator communities (Garibaldi et al 2014).

Recent studies have reported ecosystem service trade-offs in agroecosystems (Nelson et al 2009, Allan et al 2015, Sutter & Albrecht 2016). For example, intensive land use favors provisioning services (e.g. crop production) at the cost of other services (e.g. pollination). More specifically, increasing crop land at the expense of semi-natural habitat can largely reduce biodiversity in intensive agricultural landscapes (Allan et al 2014), and this may drive ecosystem service trade-offs through negative effects on ecosystem services that depend on biodiversity (Cardinale et al 2012). Thus, it may be impossible to maximize all ecosystem services simultaneously (Bateman et al 2013). These trade-offs underpin the European Commission’s Cost of Policy Inaction project (Braat and ten Brink 2008) and the land sharing *vs* land sparing debate (Green et al 2005), a framework that distinguishes between the spatial integration (land sharing) or separation (land sparing) of biodiversity conservation and crop production. A better understanding of the effects of landscape composition on crop production requires moving from the traditional single-service approach, whereby crop yield is studied individually, to a multiple-service framework (Bennett et al 2009), where crop yield and other services, such as biodiversity and pollination, are investigated simultaneously.

There is a general consensus that decreasing levels of biodiversity can reduce the magnitude and stability of ecosystem processes (Cardinale et al 2012, Tilman et al 2006). In intensively-managed agroecosystems, the decline in the diversity of pollinators associated with the loss of semi-natural habitat can alter not only the magnitude but also the temporal stability of animal pollination-dependent crop yield, especially when biodiversity is reduced to the low levels typical of many intensive agricultural areas (Garibaldi et al 2011a). This means that food security will not be achieved by high crop yields alone; agricultural practices should also target a stable provision of crop yield over time, as low crop yield stability can cause unpredictable negative impacts on food supply and farmer income (Schmidhuber and Tubiello 2007). Despite the importance of yield stability and the empirical evidence that the magnitude and stability of ecosystem services do not necessarily co-vary positively (Macfadyen et al 2011, Gagic et al 2012), there have been few studies on the stability of crop yield. These studies have generally found that yield stability decreases with agricultural intensification and crop pollination dependence (Garibaldi et al 2014, 2011a, 2011b; Deguines et al 2014), but the ecological mechanisms that drive these effects have received little attention.

In this study, we develop a model to predict changes in crop yield and biodiversity along a gradient of landscape composition (i.e. increasing proportions of semi-natural habitat) in agricultural systems. We focus on three ecosystem services, i.e. pollinator-independent crop yield (a provisioning service), crop pollination (a regulating or supporting service), and biodiversity *per se*. We assess the ecosystem service of pollination by measuring crop production resulting from animal pollination. Whether or not biodiversity is an ecosystem service in itself is a matter of debate; here, we consider biodiversity as such because it is directly associated with and drives supporting (e.g. nutrient cycling, primary production) as well as cultural services. We distinguish between two additive ecosystem services associated with total crop yield: the yield that results from wild animal pollination (hereafter crop pollination), and the yield that is independent from animal pollination (hereafter independent crop yield). This separation allows us to quantitatively vary the degree of pollination dependence of crops, in contrast to studies that only make a qualitative distinction between pollination-dependent and pollinator-independent crops (Ghazoul and Koh 2010). We analyse the expected biodiversity (i.e. species richness) and the magnitude and stability of crop pollination and independent crop yield, yielding a total of five ecosystem service components. We focus on how the relative proportion of semi-natural habitat and crop land in the agricultural landscape, and crop pollination dependence influence these five ecosystem service components. Specifically, we address two main questions: (i) What are the trade-offs between biodiversity and the magnitude and stability of crop pollination and independent crop yield in agricultural landscapes? and (ii) How do landscape composition (the relative proportion of semi-natural habitat and crop area in the agricultural landscape), and crop pollination dependence influence these trade-offs?

## Methods

### Agroecosystem model

We derive a model for crop biomass production in a spatially heterogeneous agricultural landscape that incorporates environmental and demographic stochasticity. Our model has two types of patches: crop land and semi-natural habitat. Crop land is used to grow annual crops with varying degrees of dependence on wild animal pollination, whereas semi-natural habitat shelters ‘wild’ plants and pollinators. This model represents intensively-managed agricultural systems, where crop land does not host significant levels of biodiversity, allowing spatial heterogeneity to be broadly defined by two patch types. Pollinators live and nest in semi-natural habitats, yet they move across the landscape to forage on either crops or ‘wild’ plants, or both. Crop land and semi-natural habitat are therefore linked by pollinators’ foraging movement. The three components of our model (pollinators, ‘wild’ plants, and crop yield) are represented by the following equations:

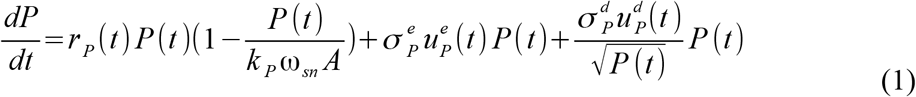

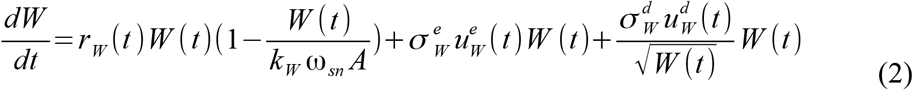

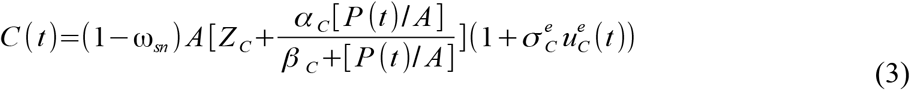

where *P* and *W* represent the maximum yearly biomass of pollinators and ‘wild’ plants, respectively. *P* does not take managed honeybees into account as they do not depend on the availability of semi-natural habitat, and they pollinate less efficiently compared to non-managed pollinators (Garibaldi et al 2013). The model does not consider within-year dynamics. *C(t)* is the amount of crop biomass produced in year *t*, i.e. annual crop yield. *C(t)* is not represented by a differential equation because crops are harvested and their dynamics do not depend on the previous state. Conversely, pollinators and wild plants are not managed and their actual values depend on previous states. *k*_*P*_ and *k*_*W*_ are the carrying capacities of pollinators and ‘wild’ plants, respectively, per unit area; *A* is the total landscape area (crop land and semi-natural habitat); ω_sn_ is the proportion of semi-natural habitat within the agricultural landscape ([1-ω_sn_] * *A* is total crop or agricultural area). The model is spatially implicit, which means that pollinators can potentially feed on all crops and ‘wild’ plants present in the agricultural landscape, irrespective of the spatial configuration of the landscape. Hence, this model describes what happens in agricultural landscapes at the scale determined by the pollinator’s foraging range (200 meters for small bee species, 25-110 meters for bumble bees, >200 meters for certain bee species (Zurbuchen et al 2010, Geib et al 2015)), which corresponds roughly to the scale of a typical arable field in Europe.

In the first two equations, *r*_*P*_ (*t*) and *r*_*w*_(*t*) are the pollinators’ and ‘wild’ plants’ per capita growth rates, and are defined as:

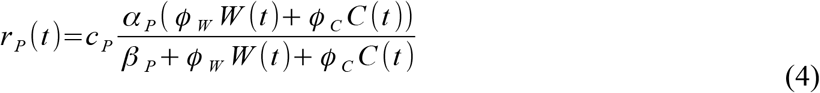

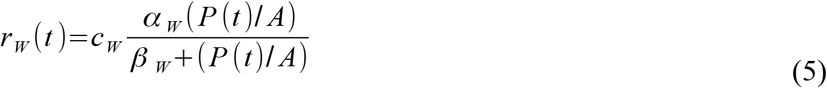

Pollinators are assumed to be generalist central-place foragers that feed on both ‘wild’ plants and crops (Kleijn et al 2015). We assume that plant and pollinator uptake of resources follows a saturating, type II functional response, where α_P_ and α_W_ are the maximum growth rates; β_P_ and β_W_ are half-saturation constants; and *c*_*P*_ and *c*_*W*_ are the conversion rates of pollinators and ‘wild’ plants, respectively, that translate the functional responses into numerical ones. For simplicity, we set conversion rates equal to unity. The pollination-dependent part of crop yield is also assumed to follow a type II functional response, where α_C_ is the maximum crop yield derived from pollination, β_C_ is the half-saturation constant of crops, and Φ_w_ and Φ_C_ are constants that convert fluxes of ‘wild’ plants and crops, respectively, to pollinator biomass. We use Φ_W_ = Φ_C_ = 1 for simplicity; to allow differences in resource quality of different crop types, we also made Φ_C_ dependent on crop pollination dependence (see below). The use of saturating functional responses is widely supported and it is consistent with several biological examples (Thebault & Fontaine 2010, Holland et al 2013, Holland 2015). A complete description of the model parameters can be found in Table 1.

**Table 1.**
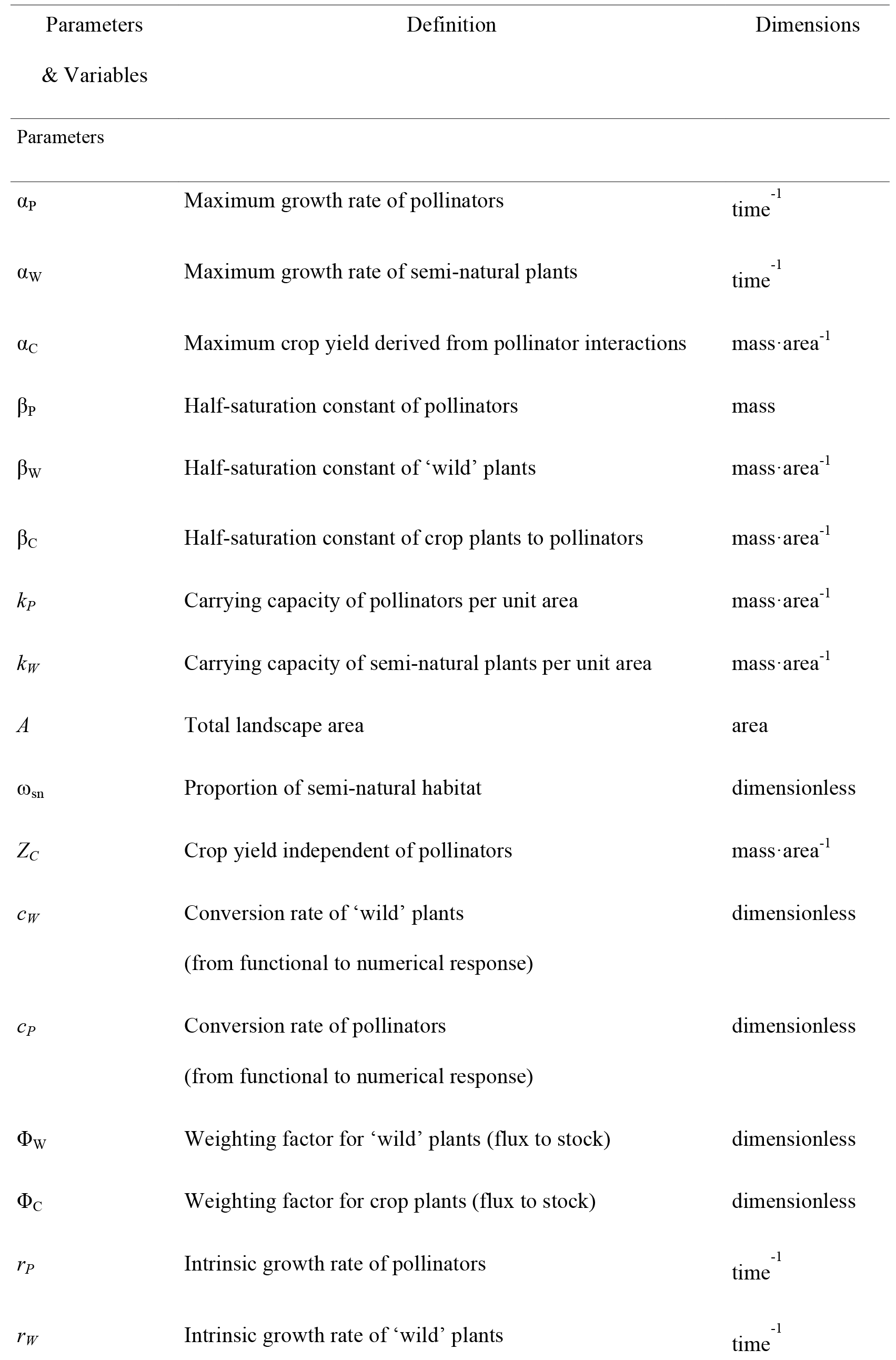
Parameters and variables of the model.

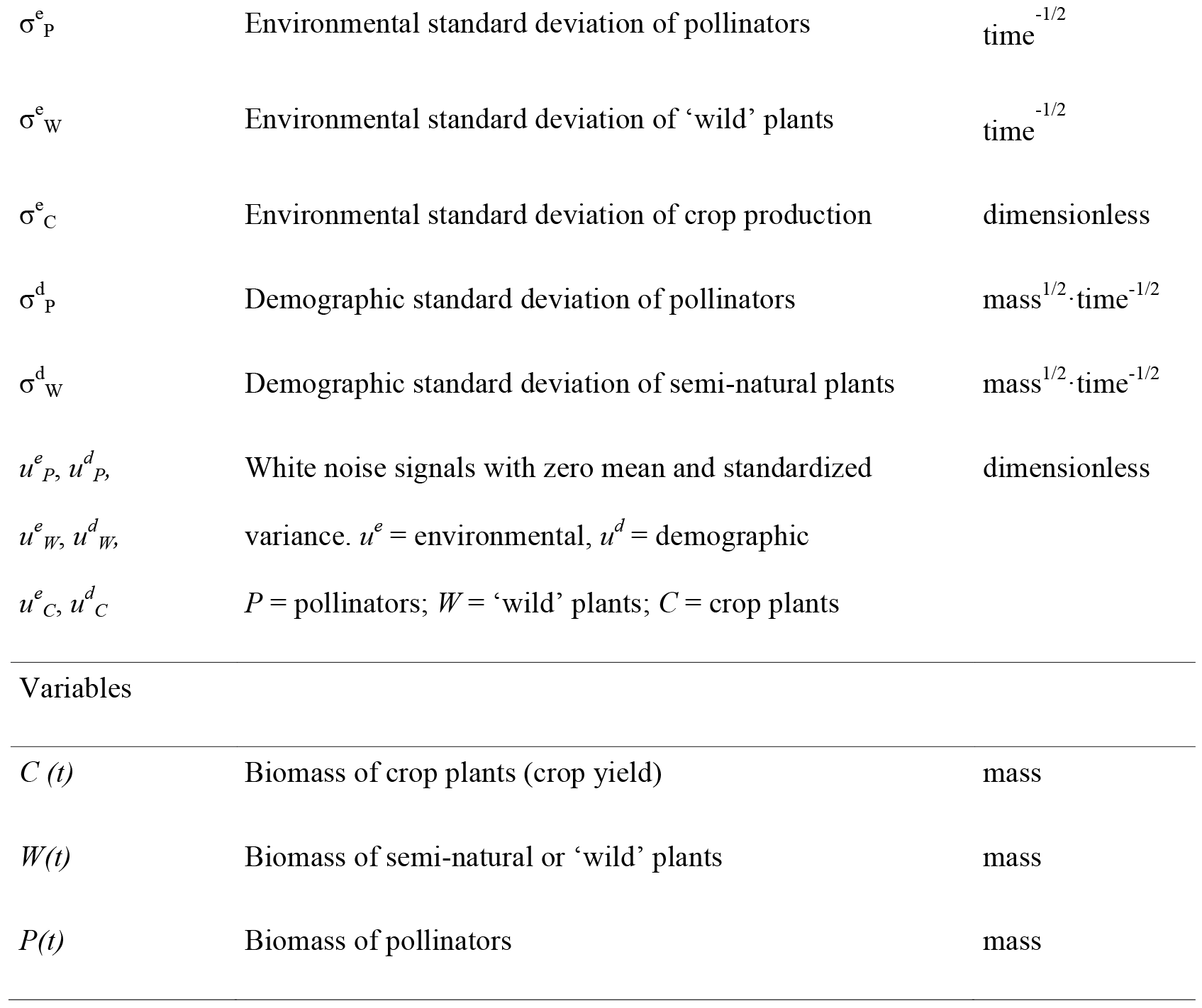

Environmental stochasticity is included through the terms σ^e^u^e^ (t), where (σ^e^)^2^ is the environmental variance of either pollinators ((σ_P_^e^)^2^), ‘wild’ plants ((σ_W_^e^)^2^) or crops ((σ_C_^e^)^2^), and u^e^(t) are random functions with zero mean and standardized variance, that can be correlated through time (a good year for plants might also be good for crops). Demographic stochasticity (σ^d^u^d^ (t)) arises from stochastic variation in individuals’ births and deaths. Because crops are sown at high densities, we assume demographic stochasticity is prevented in crops, and only affects pollinators and ‘wild’ plants. Demographic stochasticity is included in the form of the first-order normal approximation commonly used in stochastic population dynamics (Lande et al 2003), where (σ^d^)^2^ is the demographic variance of either pollinators ((σ_P_^d^)^2^) or ‘wild’ plants ((σ_W_^d^)^2^), and u^d^(t) are independent random functions with zero mean and standardized variance.

Crops differ greatly in the degree to which animal pollination contributes to yield, from pollinator-independent crops, such as obligate wind-or self-pollinated species (e.g. cereals), to fully animal-pollinated species (e.g. fruit trees, oilseed rape). Within animal-pollinated species, crops differ in their level of dependence on pollination (Klein et al 2007). In our model, *Z*_*C*_ represents the part of crop yield that is independent of animal pollination and α_C_ is the crop yield derived from pollination, and therefore we can estimate crop pollination dependence (%) as α_C_ / (α_C_ + *Z*_*C*_). If *Z*_*C*_ = 0 (α_C_ > 0), crop yield depends entirely on animal pollination; conversely, animal pollination-independent crops are defined by α_C_ = 0 (*Z*_*C*_ > 0). Most fruit and seed crops lie between these two extremes (*Z*_*C*_ > 0, α_C_ > 0). We assume there is no interaction between α_C_ and *Z*_*C*_ (Bartomeus et al 2015, Gils et al 2016).

### Mean and stability of ecosystem services

We use our model to quantify biodiversity and both the mean and the stability of independent crop yield and crop pollination, which make five ecosystem service components, in intensively-managed agricultural landscapes with varying proportions of semi-natural habitat. We assume that, at the end of each cropping season, the amount of animal pollinators, wild plants and crops reach roughly constant values in the absence of environmental and demographic stochasticity at the landscape scale, despite local year-to-year changes in those variables. This year-to-year equilibrium assumption is a reasonable first approximation to a more complex and dynamical system. We use the species-area relationship (SAR) to estimate changes in biodiversity as a function of semi-natural area. We decided to use SAR for estimating biodiversity instead of wild plant biomass or pollinator biomass, because species-biomass relationships are more variable at local/landscape scales such as the one considered here, and negative relationships have been reported (e.g. diversity-productivity) (Cardinale et al 2012). Moreover, when biodiversity is considered a cultural service, it is usually estimated as the number of species. Despite the fact that SAR is usually stronger at spatial scales larger than that of arable fields, where we might observe more variation around the average biodiversity values, it captures the expected mean biodiversity at the scale of an arable field in Europe. We estimated SAR using the conventional power law function (*S*=*c* [ω_sn_ A]^*z*^, where *S* = number of species, *c* is a constant of proportionality). Theoretical models and field data from a wide range of plant and animal taxa show that the slope, *z*, of the logarithm of species richness against the logarithm of area is roughly constant, with *z* ≈ 0.25 (Crawley and Harral 2001). Given that the equilibrium plant and pollinator biomasses are proportional to the area of semi-natural area (Appendix S5: Fig. S1), considering either species richness or biomass would yield the same qualitative results (*R*^*2*^ = 0.90; at the scale of this study, *z* can be higher (0.4 or 0.5) (Crawley and Harral 2001), yielding an even stronger correlation between the number of species and biomass). We assume that crops are harvested yearly; hence, average crop yield represents the temporal mean of the yearly averaged crop yield across the agricultural landscape. To account for the stability of independent crop yield and crop pollination, we use the inverse of temporal variability, i.e. invariability. Temporal variability is measured as the square of the temporal coefficient of variation (*CV*^*2*^) of total biomass, i.e. the ratio of the variance to the square of the mean, and is calculated in the stationary regime around the equilibrium. We use *1/CV*^*2*^ as a metric of stability (i.e. invariability) of independent crop yield and crop pollination. This measure of ecosystem stability has been used in recent empirical and experimental studies (Tilman et al 2006, Loreau and De Mazancourt 2013).

The analytical expressions for the equilibrium and variability of pollinator biomass, wild plant biomass, and crop yield are presented in Appendix S1. A summary of the equations for the five ecosystem service components can be found in Appendix S2 (Eqs. S5-S9). Whenever possible, we estimated parameter values with empirical information. In other cases, we informed parameters with commonly-assigned values found in the literature (McCann et al 2005, Thompson et al 2006, Leroux and Loreau 2008, Holland and DeAngelis 2010, Thebault and Fontaine 2010, Morales 2011, Holland et al 2013, Encinas-Viso 2014, Gounand et al 2014). For example, to determine the carrying capacity of pollinators (*k*_*P*_), we used empirical data on average numbers of individuals and body mass of wild pollinators (Bommarco et al 2012, Rollin et al 2013, Holzschuh et al 2016). For wild plants, we used empirical observations to inform their carrying capacities (*k*_*W*_) (Craven et al 2016). Also, there is information on independent crop yield that was used to determine *Z*_*C*_ (e.g. http://data.worldbank.org/). We allowed variation in α_C_ and β_C_ in order to investigate changes in the five ecosystem services components across the amount of semi-natural habitat (ω_sn_), the degree of crop pollination dependence (*Z*_*C*_/α_C_), and the crop relative requirement for pollinator densities (β_C_/*k*_*P*_). A sensitivity analysis was performed for parameter whose values could not be determined precisely or for which there was variation in their values assigned in the literature, e.g. α_C_, α_P_, *Z*_*C*_, β_C_, β_P_, *k*_*P*_ (Appendix S3). The choice of these parameters for the sensitivity analyses is also justified because they are most relevant for the estimation of equilibrium biomasses. Sensitivity analysis shows that variations in these parameter values did not change the results qualitatively. Analyses were performed in R software (R version 3.2.4, R Core Team 2016).

## Results

### Overall effects of landscape composition on ecosystem service components

Increases in the relative proportion of crop land has contrasting effects on the various ecosystem services. As expected, biodiversity increases with the proportion of semi-natural habitat, as the latter provides area for many taxonomic groups, such as wild plants and pollinators (Figure 1a). Changes in the biomasses of wild plants and pollinators with semi-natural habitat are positively correlated with changes in biodiversity (*R*^*2*^ = 0.90; Appendix S5: Fig. S1). The responses of the pollination-independent and pollination-dependent (i.e. crop pollination) components of crop yield differ strongly. Independent crop yield decreases linearly with the amount of semi-natural habitat because crop land decreases and it does not depend on pollinators (Figure 1c). In contrast, the relationship between crop pollination and the proportion of semi-natural habitat is hump-shaped (Figure 1b), as a result of the contrasting effects of semi-natural habitat on pollinators and crop land. That is, a larger amount of semi-natural habitat increases wild pollinator biomass (Appendix S5: Fig. S1b) but reduces crop land, which results in a hump-shaped relationship that is robust to changes in parameter values (Appendix S3). Total crop yield, i.e. pollination-independent plus pollination-dependent crop yields, displays a similar hump-shaped relationship, especially when crop pollination dependence is moderate to high (Appendix S5: Fig. S2). Interestingly, when measured per unit of crop land, crop yield increases with the proportion of semi-natural habitat, because of the beneficial effect of pollination (Appendix S5: Fig. S3).

**Figure 1.**
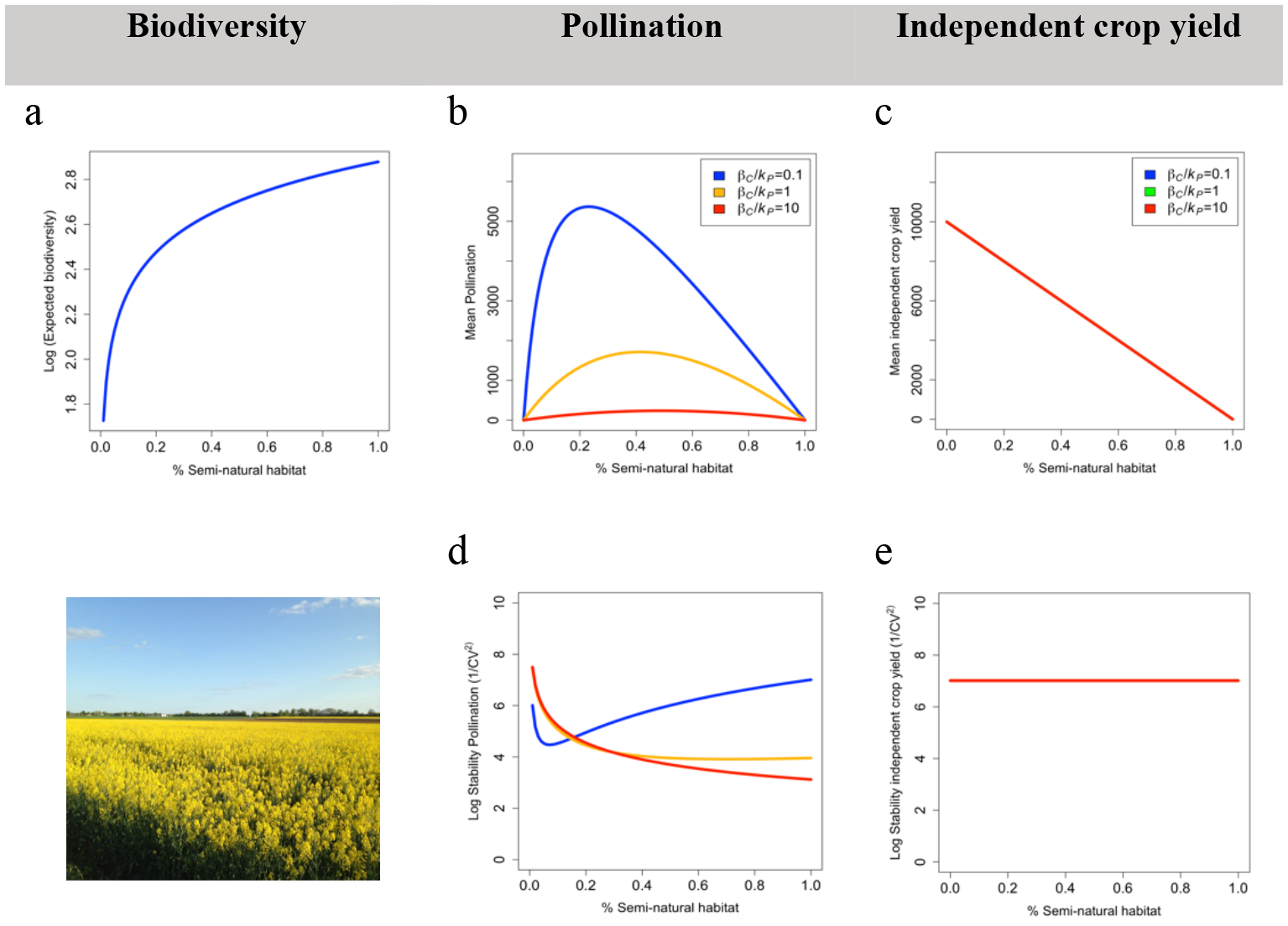
Mean and stability of five ecosystem service components in agroecosystems. This graph shows the expected biodiversity (a) and the temporal mean (c) and stability (d; log[*1/CV*^*2*^]) of independent crop yield, and crop pollination (b, d), as functions of the proportion of semi-natural habitat, for different crop relative requirement for pollinators (β_C_ /*k*_*p*_). Because β_C_ /*k*_*P*_ does not affect biodiversity and the mean/stability of independent crop yield, a single line is showed. Picture of an intensive agricultural landscape in the LSTER Zone Atelier Plaine & Val de Sèvre, France (Photo credit: Sabrina Gaba). (Parameter values: α_P_ = 0.9, β_P_ = 0.6, *A* = 10, *Z*_*C*_ = 1000, α_C_ = 1000, *k*_*W*_ = 5000, *k*_*P*_ = 0.1, σ^e^_P_ = 0.8, σ^d^_P_ = 0.1, σ^e^_C_ = 0.03, σ_C_ = 1000, Pollination dependence = 50%; Species-area relationship [*S*=*c* (ω_sn_ *A*)^z^]: c=10,
*z*=0.25)

The stability of independent crop yield does not change with semi-natural habitat (Figure 1e) because it does not rely on animal pollination. On the other hand, pollination-dependent yield does depend on animal pollinators, thus crop pollination stability strongly depends on the amount of semi-natural habitat (Figure 1d). Crop stability shows similar trends when measured at landscape scale or per unit of agricultural area.

### Role of pollination dependence and crop relative requirement for pollinators

The dependence of crop yield mean and stability on the proportion of semi-natural habitat is controlled by two *effective* parameter combinations, *Z*_*C*_ /α_*C*_ and β_C_ /*k*_*P*_ (Appendix S1). *Z*_*C*_ is the pollinator-independent component of crop yield and α_C_ is the maximum crop yield derived from pollinator interactions, so *Z*_*C*_ /α_C_ is inversely related to crop pollination dependence:

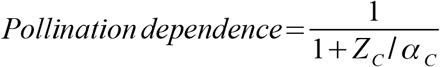

β_C_ /*k*_*P*_ is the ratio of crop half-saturation constant relative to pollinators’ carrying capacity, and it quantifies the pollinator requirement of crops relative to the availability of pollinators, i.e. crop relative requirement for pollinators. For small values of β_C_ /*k*_*P*_ (<1) crop yield saturates at lower pollinator biomass than their carrying capacity, but for large values of β_C_ /*k*_*P*_ (>1) crop yield saturates at pollinator biomasses much higher than their carrying capacities.

Biodiversity is negatively correlated with mean independent crop yield, and is unrelated to its stability (Figure 1a, c). For increasing levels of pollination dependence, both the mean and stability of total crop yield are increasingly affected by pollination and hence by the amount of semi-natural habitat (Figure 2). The position of the maximum yield along the semi-natural gradient changes with crop pollination dependence and crop relative requirement for pollinators. On one hand, for higher levels of pollination dependence crops require more pollinators and thus maximum crop yield is achieved at larger proportions of semi-natural habitat. On the other hand, high crop relative requirement for pollinators (high β_C_ / *k*_*P*_) has the dual effect of reducing mean yield and shifting maximum yield to larger amounts of semi-natural habitat. In general, high crop relative requirement for pollinators is less responsive to the amount of semi-natural habitat, because pollinator densities that will be achieved in the agricultural landscape are unlikely to fulfill crop relative requirement for pollinators (Appendix S4). Mean crop yield per unit of agricultural area increases with the proportion of semi-natural habitat (Appendix S5: Fig. S3), although it starts to show some saturation when crop relative requirement for pollinators is low. Finally we explored the effect of resource quality of different crop types and showed that these results are robust to differences in resource quality of different crop types (e.g. Φ_c_ ~ α_C_ / (α_C_ + *Z*_*C*_)) (Appendix S4).

**Figure 2.**
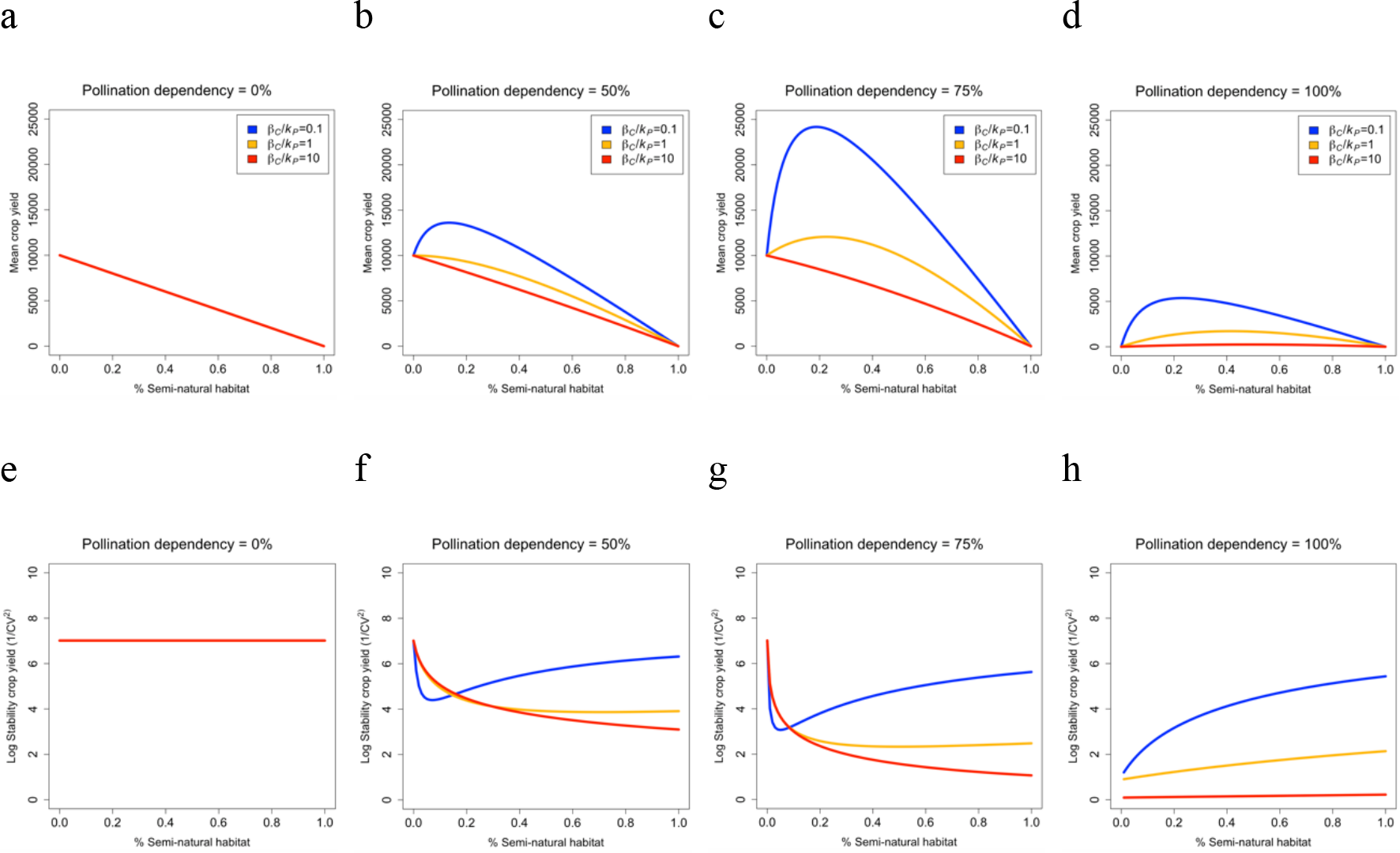
Mean and stability of total crop yield. Temporal mean and stability (log[*1/CV*^*2*^]) of total crop yield as functions of pollination dependence and crop relative requirement for pollinators. In (*a*, *e*), the three curves overlap. (Parameter values:α_P_ = 0.9, β_P_ = 0.6, *A* = *k*_*w*_ = 5000, σ^e^_P_ = 0.8, σ^d^_P_ = 0.1, σ^e^_C_ = 0.03, σ_C_ = 1000. Because *Z*_*C*_ = 1000, α_C_ is allowed to increase with higher pollination dependences; this is why mean crop yield increases with pollination dependence of crops. In a, e: α_C_ = 0 and *Z*_*C*_ = 1000. In d, h: α_C_ = 1000 and *Z*_*C*_ = 0)

In pollination-dependent crops, the stability of pollination also changes with the fraction of semi-natural habitat: it first decreases (due to the demographic and environmental stochasticity of pollinators), and then increases after a minimum fraction of semi-natural habitat has been reached (due to a drop in the response of crops to pollinator stochasticity), although this response is heavily conditioned by the crop relative requirement for pollinators (Figure 2e-h; Appendix S4). Whereas a higher pollination dependence of crops reduces pollination stability and broadens the range of stability values, crops with a lower pollination dependence are little affected by pollinator stochasticity, and yield stability is mostly determined by the environmental stochasticity of crops. Within each level of crop pollination dependence (Figure 2) the response of yield stability to semi-natural habitat is conditioned by crop relative requirement for pollinators: a low crop relative requirement for pollinators (low β_C_ /*k*_*P*_) shifts the stability valley to lower fractions of semi-natural habitat, and stability increases faster. Increasing β_C_ /*k*_*P*_ expands the region of low stability, and stability requires larger areas of semi-natural habitat to increase. When crop relative requirement for pollinators is very high (high β_C_ /*k*_*P*_), crop yield stability decreases monotonically along the full gradient of semi-natural habitat.

In sum, the contrasting effects of increasing crop land on the various ecosystem services reveal trade-offs (negatively correlated responses) and synergies (positively correlated responses) in the response of biodiversity and the mean and stability of independent crop yield and crop pollination (Figure 3). The exact shape of the ecosystem service trade-offs across the gradient of semi-natural habitat is controlled by the degree to which crops depend on pollination (*Z*_*C*_/α_C_) and by their relative requirement for pollinator densities (β_C_ /*k*_*P*_). Variations in parameter values did not change results qualitatively.

**Figure 3.**
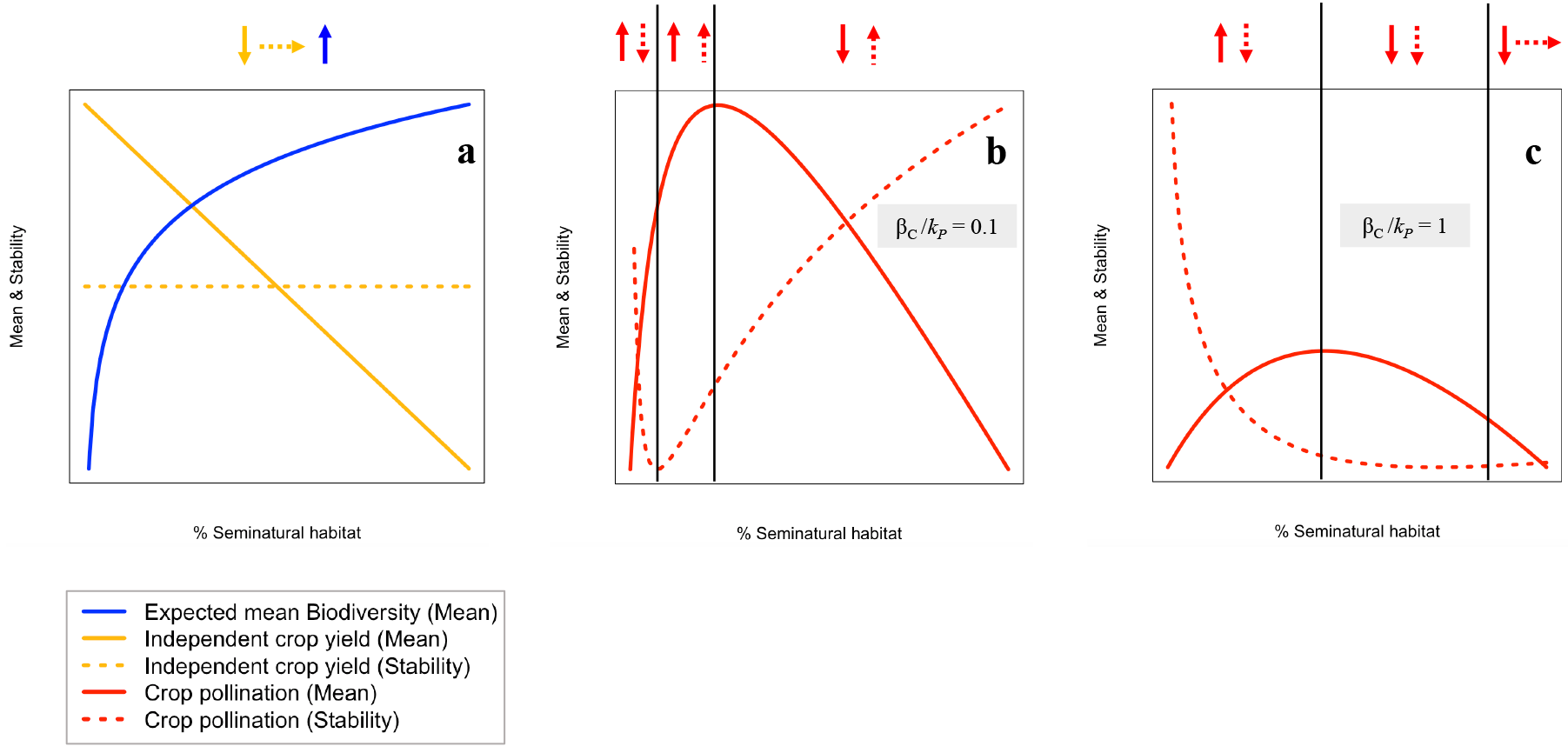
A variety of trade-offs and synergies between ecosystem service components in agroecosystems. This graph shows the expected biodiversity and the temporal mean and stability (log[*1/CV*^*2*^]) of independent crop yield (**a**), and crop pollination (**b, c**), as functions of the proportion of semi-natural habitat. The responses of the mean and stability of the three ecosystem services to increasing proportions of semi-natural habitat produce different patterns across the gradient of semi-natural habitat, from tradeoffs (negatively correlated responses: opposite arrows) to synergies (positively correlated responses: similar arrows). Independent crop yield and biodiversity (**a**) are not affected by crop relative requirement for pollinators (β_C_ /*k*_*p*_). Conversely, crop pollination mean and its stability, and therefore ecosystem service synergies and tradeoffs, are controlled by crop relative requirement for pollinators: (**b**) β_C_/*k*_*p*_= 0.1, (**c**) β_c_ /*k*_*p*_ = 1 (see main text and Supplementary Methods 1). Y axes are unit-less to make comparison between curves clearer. (Parameter values: α_P_ = 0.9, β_P_ = 0.6, *A* = 10, *Z*_*C*_ = 1000, α_c_ = 1000, *k*_*w*_= 5000, *k*_*P*_ = 0.1, σ^e^_P_ = 0.8, σ^d^_P_ 0.1. σ^e^_c_ = 0.03, α_c_ = 1000, Pollination dependence = 50%; Species-area relationship [*S*=*c* (ω_sn_*A*)^z^]: c=10, z=0.25)

## Discussion

In intensively-managed agricultural systems, increases in the amount of crop land relative to that of semi-natural habitat have major consequences for the provision of multiple ecosystem services. Our model suggests that: (1) changes in landscape composition generate a variety of synergies and trade-offs between biodiversity conservation, crop pollination and independent crop yield, (2) these trade-offs affect not only the magnitude but also the stability of these ecosystem services, and (3) the trade-offs involving crop pollination are strongly affected by the degree to which crops depend on pollination and by their relative requirement for pollinator biomass.

The loss of semi-natural habitat has contrasting effects on the three ecosystem services considered: biodiversity decreases, independent crop production increases, while pollination-dependent crop production is maximized at an intermediate proportion of semi-natural habitat. These results provide rigorous theoretical foundations for previously hypothesized functional relationships between the magnitude of ecosystem services and landscape composition (Braat and ten Brink 2008). The results further suggest that the exact shape of the hump-shaped relationship between provisioning services and semi-natural habitat is determined by the pollination dependence and the relative requirement of crops for pollinator densities (Figure 2, 3). Greater values of these two factors increase the effect of pollinator biomass on total crop yield, and thus the maximum yield is achieved at higher fractions of semi-natural habitat.

Importantly, our results suggest that landscape composition also imposes trade-offs on the stability of crop yield. These trade-offs are driven by mechanisms associated with the stochasticity of pollinators and the response of crops to that stochasticity. On the one hand, the stability of crop pollination decreases with the amount of semi-natural habitat when the latter is small because pollinator stochasticity increases. For larger proportions of semi-natural habitat, however, the response of crop yield to pollinator stochasticity drops, with varying effects on pollination stability. The decay in crop response to pollinator stochasticity is caused by the saturation of pollination-dependent crop yield to pollinator biomass (Appendix S4). Crop relative requirement for pollinators controls how fast saturation sets in and, consequently, how fast the response of crops to pollinator stochasticity drops down.

Taken together, the responses of the mean and stability of ecosystem services to landscape composition produce different patterns across the gradient of semi-natural habitat, from trade-offs (negatively correlated responses) to synergies (positively correlated responses) (Figure 3). At the landscape scale, we found a trade-off between independent mean crop yield and biodiversity, and between crop pollination and independent crop yield when semi-natural habitat is low. Conversely, at low fractions of semi-natural habitat, we observed a synergy between crop pollination and biodiversity. Such synergy between crop production and biodiversity also became apparent when considering crop production per unit of agricultural area, revealing the possibility to reconcile farmers’ profitability (at field scale) and biodiversity conservation (at landscape scale). Trade-offs and synergies can also occur within ecosystem services, e.g. crop pollination mean and its stability co-vary negatively except at low-to-intermediate amounts of semi-natural habitat. These patterns give moderate support to the intermediate landscape-complexity hypothesis (Tscharntke et al 2012), which states that the effectiveness of agro-environmental management strategies is higher in simple (1-20% non-crop area) than in either cleared (<1% non-crop area) or complex (>20% non-crop area) landscapes. For moderate-to-high levels of crop pollination dependence and high crop relative requirement for pollinators, increases in the amount of semi-natural habitat benefit biodiversity and crop pollination both in terms of average provision and stability in simple landscapes. Despite simple agricultural landscapes are often areas where cultivated crops have a low degree pollination dependency (except from species like oilseed rape and sunflower), these benefits are also larger in simple landscapes when crop yield per unit of agricultural land is considered. Surprisingly though, with a few exceptions (e.g. Duflot et al 2015), most intensively-managed agricultural landscapes show very low proportions of semi-natural habitat (<5%; Öckinger and Smith 2007, Henckel et al 2015). Additionally, increasing in the amount of semi-natural habitat benefits other services such as pest control (Sutter and Albrecht 2016). In sum, consistent with empirical observations (Pywell et al 2015, Tamburini et al 2016), the existing trade-offs and synergies suggest that moderate increases in the amount of semi-natural habitat in simple agricultural landscapes (1-20% non-crop area) allow ecosystem services essential for crop production to be maintained, which in turn increases the magnitude and stability of crop yield.

Our findings are also consistent with recent studies suggesting that the interaction between agricultural intensification and the level of pollination dependence of crops determines the stability of crop production at large spatial scales. For instance, using an intensification index that includes the amount of semi-natural habitat in agroecosystems, a recent study found that the stability of the yield of the 54 major crops in France decreases in more intensive agriculture, and that this reduction is more pronounced for higher crop pollination dependence (Deguines et al 2014). Similarly, long-term data from FAO suggest that a greater pollination dependence of crops leads to lower and less stable crop yields (Garibaldi et al 2011a). By considering multiple inter-related ecosystem services simultaneously, our results add a mechanistic understanding of these ecosystem service trade-offs in intensively-managed agroecosystems.

The trade-offs in ecosystem service provision revealed by our model have two major implications for the management of intensive agricultural systems. First, the effects of biodiversity loss on crop production that result from agricultural intensification depend on the level of pollination dependence of crops. Whereas in pollinator-independent agriculture reductions in biodiversity and crop pollination have no effect on provisioning services (crop production), for pollination-dependent agriculture crop production relies on biodiversity (e.g. wild plants provide foraging, nesting and refuge for pollinators), and the trade-off between biodiversity conservation and crop production is mediated by biodiversity loss. Such reduction in biodiversity reduces the delivery of regulating services, and this has a direct negative effect not only on mean yield but also on its stability. Secondly, our results suggest that simultaneously maximizing crop yield mean and stability is often impossible for pollination-dependent crops, and therefore, management strategies that focus on maximising mean yield will overlook its stability. Specifically, enhancing crop yield by increasing crop land would be counterproductive for pollination-dependent crops, at least below a threshold of semi-natural habitat. There is, however, a notable exception to this: maximization of crop yield mean (both at the landscape scale and per unit of agricultural area) and crop yield stability can be achieved at 20-40% of semi-natural habitat when crops show intermediate-to-high degrees of animal pollination dependence and crop relative requirement for pollinators is low.

The yield mean and stability of crops with greater pollinator dependence has continuously decreased from 1961 to 2008 (Garibaldi et al 2011a). This suggests that the relative requirement for pollinators of many world crops is high, as pesticide use has diminished the carrying capacity of pollinators in semi-natural habitat during the same period of time (Goulson et al 2015). To compensate for low crop yields agricultural policies have promoted land cultivation of pollination-dependent crops and the use of managed honeybee colonies, which are not affected by semi-natural habitat. However, these measures reduce the amount of semi-natural area and honeybees cannot compensate for the pollination services of non-managed, wild pollinators (Aizen et al 2009, Garibaldi et al 2013). Our model suggests that an alternative to agricultural intensification consists in diminishing crop relative requirement for pollinators with practices that increase the carrying capacity of pollinators in semi-natural habitat, such as higher farmland heterogeneity and floral assemblages, increasing nesting opportunities, and reductions in the use of synthetic pesticides (Garibaldi et al 2014). These measures may not only increase mean crop yield at the landscape scale or per unit of agricultural area, but also its stability.

Our model has several limitations. For example, our model and the observed trade-offs between biodiversity and crop yield refer to intensively-managed agricultural systems, where crop land does not host important biodiversity levels; however, these trade-offs are not necessarily similar in non-intensive agricultural systems where biodiversity can moderately thrive within crop land (Clough et al 2011). Second, the species-area relationship is stronger at spatial scales larger than that of arable fields, where we might expect more variation around the expected biodiversity values; yet, our simple model captures the expected mean biodiversity at the scale of an arable field in Europe. Besides, the observation that biodiversity loss has either none (stability) or positive (mean) effects on independent crop yield is based on the species-area relationship; these effects are likely to differ if taxonomic groups responsible for other ecosystem services, i.e. pest control, are more specifically included. Also, our model is spatially-implicit, and does not consider the effects of the spatial configuration of semi-natural habitat (Garibaldi et al 2011b, Mitchell et al 2015); future studies should consider space explicitly, as the spatial distribution of semi-natural habitat within the agricultural landscape determines the ecosystem service flows between semi-natural habitat and crop land, including pollination (Brosi et al 2008, Keitt 2009, Serna-Chavez et al 2014). Finally, we find that the amount of semi-natural habitat has no effect on the stability of independent crop yield. This may change, however, if environmental stochasticity of crops increases with decreasing amounts of semi-natural habitat, as suggested by studies linking semi-natural habitat to climate regulation, natural hazard regulation and water flow regulation services (Harrison et al 2010). Despite these limitations, our model is a very useful first step as it successfully reproduces the results of recent empirical studies on the stability of pollination-dependent crop yield and it provides a mechanistic understanding of the trade-offs that are relevant in intensively-managed agroecosystems.

## Conclusions

Although historically the demand for increased crop production has been satisfied by agricultural practices that promote land conversion to crop land and improvements in crop yield (e.g. fertilizers, pesticides, selection of high-yield crop strains), the benefits of this approach have started to be challenged. The present study sheds new light on this debate. Our model suggests that landscape composition imposes trade-offs on several ecosystem services in intensively-managed agroecosystems. These trade-offs not only affect the mean production of crops, but also their temporal stability, in such a way that high and stable crop yields are not necessarily associated. This suggests that an approach that simultaneously considers the magnitude and stability of multiple ecosystem services is needed to understand and better manage agricultural systems. In order to develop a more efficient agriculture and ensure food security, it is essential to understand the mechanisms driving the trade-offs between multiple ecosystem services.

## Acknowledgements

DM was funded by the EU and INRA in the framework of the Marie-Curie FP7 COFUND People Program, through the award of an AgreenSkills/AgreenSkills+ fellowship. This work was supported by the TULIP Laboratory of Excellence (ANR-10-LABX-41), by the ANR AGROBIOSE (ANR-13-AGRO-0001), ERANET ECODEAL, and by the BIOSTASES Advanced Grant, funded by the European Research Council under the European Union’s Horizon 2020 research and innovation program (grant agreement No 666971). The authors thank the Centre for Biodiversity Theory and Modelling (CBTM) laboratory members for helpful discussion

